# Production of multiple bacteriocins, including the novel bacteriocin gassericin M, by *Lactobacillus gasseri* LM19, a strain isolated from human milk

**DOI:** 10.1101/841254

**Authors:** Enriqueta Garcia-Gutierrez, Paula M. O’Connor, Ian J. Colquhoun, Natalia M. Vior, Juan Miguel Rodríguez, Melinda J. Mayer, Paul D. Cotter, Arjan Narbad

## Abstract

Bacteriocins are antimicrobial peptides produced by bacteria and their production by health-promoting microbes is regarded as a desirable probiotic trait. We found that *Lactobacillus gasseri* LM19, a strain isolated from human milk, exhibits antagonistic activity against different enteropathogens and produces several bacteriocins, including a novel bacteriocin, gassericin M. These bacteriocins were purified from culture and synthesised to investigate their activity and potential synergy. *L. gasseri* LM19 was tested in a complex environment mimicking human colon conditions where it not only survived but expressed the seven bacteriocin genes and produced short chain fatty acids. Metagenomic analysis of these *in vitro* colon cultures showed that co-inoculation of *L. gasseri* LM19 with *Clostridium perfringens* gave profiles with more similarity to controls than to vessels inoculated with *C. perfringens* alone. This makes *L. gasseri* LM19 an interesting candidate for further study for maintaining homeostasis in the gut environment.

## Introduction

Beneficial bacteria have consistently been harnessed throughout human history. Most recently, the rise of antimicrobial resistance among pathogens, a greater demand for healthy foods and an increasing appreciation of the importance of the human gut microbiota have brought attention back to natural sources of new antimicrobials, food preservatives and probiotics. The search for natural antimicrobials can involve a variety of approaches (Lewis, 2013), including taking advantage of the fact that, in nature, bacteria from a specific environmental niche are able to compete against other bacteria from the same niche in a variety of ways (Czárán *et al.*, 2002; Kelsic *et al.*, 2015). Such bacterial antagonism can be through non-specific strategies, like the production of organic acids. Some organic acids, particularly the short-chain fatty acids (SCFA), acetate, propionate, and butyrate, are produced in millimolar quantities in the gastrointestinal (GI) tracts of animals and humans and, in addition to their antagonistic activities, confer other health benefits (LeBlanc *et al.*, 2017; Singh *et al.*, 2018). Target-specific antagonistic activities can be provided by compounds such as bacteriocins (Garcia-Gutierrez *et al.*, 2018), a heterogeneous group of ribosomally-synthesised peptides that represent a potential alternative to traditional antibiotics because of their frequent low toxicity, high potency, ability to be bioengineered, low likelihood of resistance development and the possibility of being produced *in situ* by probiotics (Cotter *et al.*, 2013; Field *et al.*, 2015; Hegarty *et al.*, 2016).

*Lactobacillus* spp. are members of the lactic acid bacteria (LAB) and contribute to the production of many fermented foods, as well as being important components of the human gut microbiota. *Lactobacillus* and other LAB are considered an important source of antimicrobial peptides (Collins *et al.*, 2017). *Lactobacillus gasseri* is one of six species which previously comprised the *L. acidophilus* complex (Fujisawa *et al.*, 1992; Sarmiento-Rubiano *et al.*, 2010). These species are considered ecologically and commercially important and have been extensively studied, frequently revealing antimicrobial and other probiotic properties (Abramov *et al.*, 2014; Karska-Wysocki *et al.*, 2010; Kim *et al.*, 2007; Selle and Klaenhammer, 2013; Yamano *et al.*, 2007). *L. gasseri* has been divided in two subgroups on the basis of average nucleotide identity (ANI) (Tada *et al.*, 2017) and strains have been previously isolated from the gut of animals and humans, vaginal tract, human milk and oral cavity. Strains of *L. gasseri* have been found to produce bacteriocins, frequently referred to as gassericins, corresponding to different classes. Gassericin A is a cyclic class IIc bacteriocin produced by *L. gasseri* LA39 that was isolated and purified from a human infant faecal sample (Kawai *et al.*, 1994; Pandey *et al.*, 2013). Gassericins B1, B2, B3 and B4 were isolated from vaginal isolate *L. gasseri* JCM 2124 with B1 and B3 being identical to the α and β peptides of the two-component bacteriocin acidocin J1132 from *Lactobacillus acidophilus* and B2 and B4 suggested to be modified forms of B3 (Tahara *et al.*, 1997). Production of gassericin T (GasT) was first reported in *L. gasseri* SBT 2055, a strain isolated from adult human faeces (Kawai *et al.*, 2000). The amino acid sequence of GasT shows high similarity to one of the peptides (LafA) of the two-component lactacin F family produced by *Lactobacillus johnsonii* VPI11088 (Kawai *et al.*, 2000). Along with the LafA peptide, *L. johnsonii* produces another hydrophobic peptide, LafX, which was highly similar to lactobin A and the predicted product of *gatX* found in the operon of *L. gasseri* SBT 2055 (Kawai *et al.*, 2000). GatX was ultimately detected and its antimicrobial activity was confirmed (Mavrič *et al.*, 2014) and the corresponding bacteriocin has been assigned to class IIb. This was considered an important finding, because in some instances where two peptides are involved, one of the peptides is usually described as active and the other as the complementary factor (cf) without antimicrobial activity, based on similarities with lactacin F complementary component. Acidocins LF221A and LF221B were isolated from *L. acidophilus* LF221 (later renamed *L. gasseri* LF221), a strain isolated from infant faeces (Bogovic-Matijasic *et al.*, 1998), but the suggested assignment to the two-peptide bacteriocin group (Class IIb) has not been established experimentally (Maldonado-Barragán *et al.*, 2016). *L. gasseri* K7 was also isolated from the faeces of a breast-fed baby and two two-peptide bacteriocin-encoding operons were found in its genome (Zorič Peternel *et al.*, 2010). These potential peptides shared a high homology to acidocins LF221A and LF221B and gassericin T peptides, respectively (Mavrič *et al.*, 2014). Isolation and purification of gassericin E from *L. gasseri* EV1461 isolated from the vagina of a healthy woman has been reported (Maldonado-Barragán *et al.*, 2016). Gassericin E exhibits high similarity to gassericin T, differing by only by one amino acid residue across the mature peptide (Maldonado-Barragán *et al.*, 2016). Interestingly, the gassericin E operon also presents a putative bacteriocin-encoding gene, *gaeX*, whose product shares 100% identity with GatX of *L. gasseri* SBT 2055 and gassericin K7 B of *L. gasseri* K7 (Maldonado-Barragán *et al.*, 2016). Finally, genes encoding gassericin T (GatA and GatX) and the novel gassericin S, with similarity to acidocin LF221A (GasA and GasX), were all found in the genome of *L. gasseri* LA327, isolated from human large intestine tissue (Kasuga *et al.*, 2019). Kasuga et al demonstrated the synergistic activity between the two components of gassericin T, and those of gassericin S (Kasuga *et al.*, 2019). However, they could not demonstrate synergistic activity between gassericin S and gassericin T when they mixed the four peptides together.

Here we report a new *L. gasseri* strain, LM19, isolated from human milk, which possesses three bacteriocin clusters in its genome, including one encoding a novel bacteriocin, designated gassericin M. Genes with homology with the paired peptides of gassericin T and gassericin S were also found in its genome. All six peptides were purified and tested for antimicrobial activity. We also demonstrated that LM19 survives, expresses all the bacteriocin genes and produces SCFA in detectable amounts in a complex faecal environment mimicking colon conditions, and that it can help to maintain the composition of the microbiome in the presence of the pathogen *Clostridium perfringens*.

## Methods

### Isolation and whole genome sequencing of *L. gasseri* LM19

*L. gasseri* LM19 was originally isolated from breast milk on MRS agar (Oxoid) at 37°C and has been deposited in the National Collection of Industrial, Food and Marine Bacteria (NCIMB) with the accession number NCIMB 15251. Whole genome sequence was provided by MicrobesNG (Birmingham, UK) using Illumina® HiSeq and a 250 bp paired end protocol. Genome coverage was 30x. Reads were trimmed using Trimmomatic 0.30 with a sliding window quality cutoff of Q15 (Bolger *et al.*, 2014) and the quality was assessed using software Samtools (Li *et al.*, 2009), BedTools (Quinlan and Hall, 2010) and BWA mem (Li and Durbin, 2010). SPAdes 3.7 (Bankevich *et al.*, 2012) was used for *de novo* assembly and annotation was performed using Prokka 1.11 (Seemann, 2014).

### Bioassay-based screening for antimicrobial activity

For antimicrobial overlay assays, 5 µl aliquots of *L. gasseri* LM19 overnight cultures were spotted onto agar plates (containing 2% w/v NaHCO_3_ to counteract inhibition from lactic acid) and incubated for up to 48 h. Bacterial spots were exposed to UV light for 15 min before being covered with 5 ml soft agar (0.7%) cooled to <50°C and inoculated with 100-200 µl of an overnight culture of an indicator strain. Overlaid plates were incubated overnight at the appropriate conditions for the indicator strain. Antimicrobial activity was considered positive if a zone of inhibition was seen (Balouiri *et al.*, 2016). For cross streak assays, *L. gasseri* LM19 was streaked onto an agar plate containing 2% NaHCO_3_ and incubated to allow growth. Streaks were exposed to UV light for 15 min and cross-streaked with different indicator strains. For drop tests, indicator strains were cultured overnight and diluted 1:100 in phosphate buffer saline (PBS). 100 µl was spread onto agar plates to produce a lawn. 10 µl of cell free supernatants of *L. gasseri* LM19 cultures, centrifuged at 16,000 × *g* for 2 min and filtered through a 0.22 µm filter (Millipore, UK) were spotted onto the lawn. For filter disc tests, the drop test method was followed but supernatants were spotted onto a 3MM Whatman filter disc that was placed onto the bacterial lawn then plates were kept at 4°C for 2 h to allow diffusion through the agar. All plates were incubated overnight in appropriate conditions for the indicator strains. For well-diffusion assays, agar plates were poured containing 1 ml overnight culture of the indicator strain. 50 µl of cell-free *L. gasseri* LM19 bacterial supernatants were placed in wells made with a cork borer; plates were kept at 4°C for 2 h to allow diffusion and incubated overnight. Inhibitory activity was assessed by measuring the radius of inhibition (mm).

To assess antimicrobial activity against *Campylobacter jejuni*, Skirrow plates (Oxoid) were inoculated with 50 μl of a *C. jejuni* stock in 40% glycerol and incubated overnight at 37°C in microaerobic conditions (85% N_2_, 5% O_2_, 10% CO_2_) in a MACS-MG-1000 controlled atmosphere cabinet (Don Whitley Scientific, UK). The following day, cells grown on the plate were resuspended in 2 ml PBS and a dilution of final optical density at 600 nm (OD_600_) = 1 was prepared in PBS. 5 ml Brucella/agar mix (1.5 g agar in 100 ml of Brucella broth (Oxoid, UK) with 0.01% triphenyl tetrazolium chloride [TTC]) were added to 1 ml cell aliquots and poured onto a fresh Brucella plate. Filter discs placed onto the agar were spotted with 10 µl *L. gasseri* LM19 cell free supernatants. 10% hydrogen peroxide was used as a positive control.

Bacterial strains used were obtained from culture collections (ATCC, American Type Culture Collection; DSMZ, Deutsche Sammlung von Mikroorganismen und Zellkulturen; NCTC, National Collection of Type Cultures) or were from in-house collections. Strains and culture conditions were: *L. gasseri* LM19 (MRS, 37°C, anaerobic, static), *Salmonella enterica* LT2 (LB, 37°C, anaerobic, static), *Escherichia coli* ATCC 25922 (LB, 37°C, anaerobic, static), *Cronobacter sakazakii* DSMZ 4485 (BHI, 37°C, anaerobic, static), *Clostridium perfringens* NCTC 3110 (BHI, 37°C, anaerobic, static), *Listeria innocua* NCTC 11288 (BHI, 37°C, anaerobic, static), *Lactobacillus delbrueckii* subsp. *bulgaricus* (*L. bulgaricus*) 5583 (MRS, 37°C, anaerobic, static), *L. bulgaricus* LMG 6901 (MRS, 37°C, aerobic, static), *Campylobacter jejuni* NCTC 11168 (Brucella, 37°C, microaerobic, static), and *Micrococcus luteus* FI10640 (MRS, 37°C, aerobic, static). Media was sourced from Oxoid.

### In silico identification of bacteriocin gene clusters

The *L. gasseri* LM19 genome was analysed with software to identify putative bacteriocin clusters: BAGEL 3 and BAGEL 4 (van Heel *et al.*, 2013) to target bacteriocin clusters and antiSMASH to target secondary metabolites (Weber *et al.*, 2015). The assembly was also annotated with RAST (Rapid Annotation using Subsystem Technology) and visualised with SEED (Brettin *et al.*, 2015). Genome data was visualised using Artemis (Carver *et al.*, 2012). DNA and amino acid sequences identified as putative bacteriocin genes and proteins were analysed using BLAST (Altschul, 1990) to assess their relationships with other peptides using default parameters. Geneious Tree Builder v11.1 (Biomatter, New Zealand) was used to compare the gassericins.

### Detection and purification of antimicrobial peptides

*L. gasseri* LM19 was grown anaerobically at 37°C in 2 l MRS broth for 24-48 h. The culture was centrifuged (8,000 × *g*, 20 min, 10°C) to separate cells from supernatant, and both cells and supernatant were analysed independently. The cell pellet was resuspended in 400 ml 70% propan-2-ol, 0.1% trifluoroacetic acid (TFA – ‘IPA’) using a stirrer for 3-4 h at room temperature, centrifuged again and the supernatant retained for further purification and activity testing by drop test using *L. bulgaricus* LMG 6901 as an indicator strain. IPA was removed from this extract by rotary evaporation until the sample volume was 120 ml, and it was applied to a 2g 12 ml Strata® C_18_-E solid-phase extraction (C18-SPE) column (Phenomenex, UK), pre-equilibrated with methanol and water following manufacturer’s instructions. The column was washed with 20 ml of 30% ethanol and 20 ml of 30% acetonitrile and the active fraction was eluted with 30 ml of IPA. The IPA was removed from the C18 SPE IPA eluate and 4 ml aliquots of sample applied to a semi preparative Jupiter C5 Reversed Phase HPLC column (10 x 250 mm, 10 μm, 300Å) (Phenomenex, Cheshire, UK) (HPLC run I) running a 30-70% acetonitrile 0.1% formic acid (FA) gradient over 95 minutes where buffer A is 0.1% FA and buffer B is 100% acetonitrile 0.1% FA. Flow rate was 2.5 ml/min and fractions were collected at 1 min intervals. The fractions were further analysed by matrix assisted laser deionisation-time of flight-mass spectrometry (MALDI-TOF-MS; Axima TOF^2^ MALDI-TOF mass spectrometer in positive-ion reflectron mode, Shimadzu Biotech, UK) to determine the masses of the potential peptides. For purification from the cell-free supernatant, the supernatant was applied to an Econo-column (BioRad, UK) containing 60 g Amberlite XAD 16N. The column was washed with 400 ml 35% ethanol followed by 400 ml 30% acetonitrile and antimicrobial activity eluted with 450 ml IPA. The IPA was removed from the XAD IPA eluate by rotary evaporation until the sample volume was 145 ml and it was then applied to a 5 g 20 ml C18-SPE column pre-equilibrated with methanol and water following manufacturer’s instructions. The column was washed with 30 ml 30% ethanol followed by 30 ml 30% acetonitrile and antimicrobial activity eluted with 30 ml IPA and fractionated by semi-preparative reversed phase HPLC as before. To increase purity, some HPLC fractions were reapplied to the C5 semi prep column, running shallower gradients. Specifically, 30-40% acetonitrile 0.1% FA gradient over 95 min for GamX and Bact_2, 30-45% gradient for GamA, and 35-65% gradient for Bact_1, GamM and GamY. Additionally, the six peptides were synthesised using microwave-assisted solid phase peptide synthesis (MW-SPPS) performed on a Liberty Blue microwave peptide synthesizer (CEM Corporation, USA). GamA and GamM were synthesised on a H-Lys(BOC)-HMPB)-ChemMatrix® resin, GamX was synthesised on H-Asn(Trt)-HMPB-ChemMatrix® resin, Bact_1 and Bact_2 on H-Arg(PBF)-HMPB-ChemMatrix® resin and GamY on Fmoc-Phe-Wang (Novobiochem®, Germany) resin. Crude peptide was purified using RP-HPLC on a Semi Preparative Vydac C4 (10 x 250mm, 5µ, 300Å) column (Grace, USA) running acetonitrile-0.1% TFA gradients specific to the peptide of interest. Fractions containing the desired molecular mass were identified using MALDI-TOF-MS on an Axima TOF2 MALDI TOF mass spectrometer and were pooled and lyophilized on a Genevac HT 4X lyophilizer (Genevac Ltd., UK). All naturally produced peptides and synthetic peptides, from HPLC runs, were assayed by well-diffusion assay using *L. bulgaricus* DPC6091 to compare activity and to assess synergistic activity among them.

### Transformation of *L. gasseri* LM19

Electro-competent cells of *L. gasseri* LM19 were made based on the method described previously (Holo and Nes, 1989). Competent cells were resuspended in 2.25 ml 10% glycerol/ 0.5 M sucrose, aliquoted in volumes of 40 µl and either used immediately or frozen on dry ice. 500 ng of plasmid pUK200 (Wegmann *et al.*, 1999) were added to 40 µl of electro-competent cells. The mixture was incubated for 1 min on ice and transferred to a pre-chilled electroporation cuvette (Geneflow Limited, UK). A pulse of 1500 V, 800 Ω and 25 µF was applied using a BioRad electroporator. 450 µl of pre-chilled MRS/ 20 mM MgCl_2_/ 2 mM CaCl_2_ were added to the cuvette and the mixture transferred to a chilled 2 ml tube and incubated for 2 h at 37°C. Aliquots were plated on MRS with 7.5 μg/ml chloramphenicol and incubated overnight at 37°C. Transformants were confirmed by colony PCR using Go Taq G2 polymerase (Promega) and primers p181 (5’-GCGAAGATAACAGTGACTCTA-3’ and p54 (5’-CGGCTCTGATTAAATTCTGAAG-3’) (Sigma, UK).

### Fermentation studies

*L. gasseri* LM19 was inoculated at 1% in 20 ml of prepared in-house MRS without glucose (10 g/l trypticase peptone (Difco, UK), 2.5 g/l yeast extract (Difco, UK), 3 g/l K_2_HPO_4_ (Sigma, UK), 3 g/l KH_2_PO_4_ (Sigma, UK), 2 g/l tri-ammonium citrate (Sigma, UK), 0.2 g/l pyruvic acid (Sigma, UK), 0.3 g/l cysteine-HCl (Sigma, UK), 0.575 g/l MgSO_4_·7H_2_O (Sigma, UK), 0.12 g/l MnSO_4_·7H_2_O (Sigma, UK), 0.034 g/l FeSO_4_·7H O (Sigma, UK) and 1 ml Tween 80 (Sigma, UK)), or batch model media, prepared as described previously (Parmanand *et al.*, 2019). The pH was adjusted to 6.8 in both media and filter sterilized carbohydrate source supplementation (glucose, lactose, galactose, inulin, starch or pectin [Sigma, UK]) was added at 2% after autoclaving. Fermentations were incubated in anaerobic conditions at 37°C over 48 h, conducted in triplicate and 2 ml of each sample were collected at 24 h and 48 h. 1 ml was used for enumeration by plate count, pH measurement using a pH-000-85282 probe (Unisense, Denmark) and, once filter sterilized, antimicrobial activity using a well diffusion assay; the other was centrifuged at 16,000 x g and the cell-free supernatant stored at −20°C for further analysis.

### *In vitro* colonic batch model fermentation

Fermentations to simulate human colon conditions were performed as described previously (Parmanand *et al.*, 2019). A faecal dilution was prepared with 10 g of fresh faecal sample diluted 1:10 in PBS, homogenised using a Circulator 400 stomacher bag (Seward, UK) in a Stomacher 400 circulator (Seward, UK) for two cycles of 45 s at 230 rpm and then transferred to pre-reduced batch model media in a proportion 1:10, to a final volume of 150 ml, in 270 ml water-jacketed vessels (Soham Scientific, UK). The temperature was maintained at 37° C by a circulating water bath. Batch model media was prepared as stated before, and 1% glucose was added as a carbohydrate source. Cultures were stirred, anaerobic conditions were maintained with oxygen-free nitrogen and pH maintained between 6.6-7.0 by adding 1M NaOH or 1M HCl with automated pH controllers (Electrolab Ltd, UK) (Parmanand *et al.*, 2019). Overnight cultures of *L. gasseri* LM19 pUK200 and *C. perfringens* NCTC 3110 were added to the vessels at 1% each. 6 ml samples were extracted at 0 h, 4 h, 8 h, 24 h and 48 h for DNA and RNA extractions, SCFA analysis and enumeration of *L. gasseri* LM19 pUK200 by plate count on MRS supplemented with 7.5 μg/ml chloramphenicol. Experiments were carried out in triplicate using three different faecal donors.

To test *L. gasseri* LM19 bacteriocin gene expression in faecal samples, 3 ml of each aliquot at different time points was treated for RNA extraction, cleaning and cDNA synthesis. Briefly, each sample was mixed with two volumes of RNAlater (Sigma Aldrich, UK) and centrifuged for 10 min at 18,000 × *g* at 4°C. The supernatant was discarded, and pellets stored at −80°C until extraction. Extraction was performed using the Qiagen RNeasy extraction kit with minor modifications. Pellets were resuspended in 1 ml RLT buffer provided in the kit, complemented with 10 µl of β-mercaptoethanol (Millipore, UK) and transferred to lysing matrix E tubes (MP Biomedicals LLC, France). Samples were lysed in a FastPrep-24 homogeniser (MP Biomedicals) by applying 2 pulses of 30 s and intensity 6.0 with an interval of 1 min on ice between each pulse. Samples were centrifuged for 10 min at 17,000 × *g* and the supernatant transferred to clean 15 ml tubes and mixed with an equal volume of 70% ethanol. 70% of the mixture, including any precipitate, was transferred to spin tubes and centrifuged at 8,000 × *g* for 1 min and following steps were as the manufacturer’s instructions. The RNA was eluted in 100 µl RNase-free water and quantified by NanoDrop 2000 (Thermo Scientific, UK). DNase treatment was performed using the Turbo DNA-free™ kit (Invitrogen, UK) following the manufacturer’s protocol. cDNA synthesis was carried out using the QuantiTect® Reverse Transcription Kit (Qiagen, UK) using 100 ng RNA per reaction. Reverse transcription was conducted according to manufacturer’s recommendations and a control reaction replacing the reverse transcriptase with water was set up at the same time. The presence or absence of *L. gasseri* LM19 bacteriocin genes was confirmed by PCR (Treven *et al.*, 2013). Primers (Table 1) were designed using Primer 3 (v. 0.4.0). The presence of secondary structures and dimers were tested with Netprimer (Premier Biosoft) and primers were tested using genomic DNA from *C. perfringens* NCTC 3110, extracted using the genomic tip-20 and genomic buffer set kits (Qiagen, Germany), to confirm their specificity. Thermal cycling was performed using a Verity 96 well Thermal Cycler (Applied Biosystems) using GoTaq G2 DNA polymerase (Promega) according to manufacturer’s instructions. Primers used are summarised in Table 1 and cycle conditions were the same as for qRT-PCR. dNTPs were provided by Bioline. PCR products were visualized using 2% agarose gels. qRT-PCR was performed using 384-well plates (4titude Ltd, UK) in the ViiA™ 7 System (Applied Biosystems, UK) with the SensiFAST™ SYBR® No-ROX Kit (Bioline, USA). Reaction mix composition was, for a final volume of 6 µl, 0.6 µl of cDNA template, 3 µl of 2x SensiFAST SYBR® No-ROX mix, 0.24 µl of each primer (10 µM stock) and 1.92 µl of water. Reaction conditions were set up at 20 sec at 95°C, 40 cycles of 1 sec at 95°C, 20 sec at 60°C and 15 sec at 95°C, and a melt curve of 15 min at 65°C. Reactions were set up in duplicate and controls for primers and no transcriptase samples were set in each run. Baseline for change was established at 2X upregulation or downregulation.

**Table 1.**
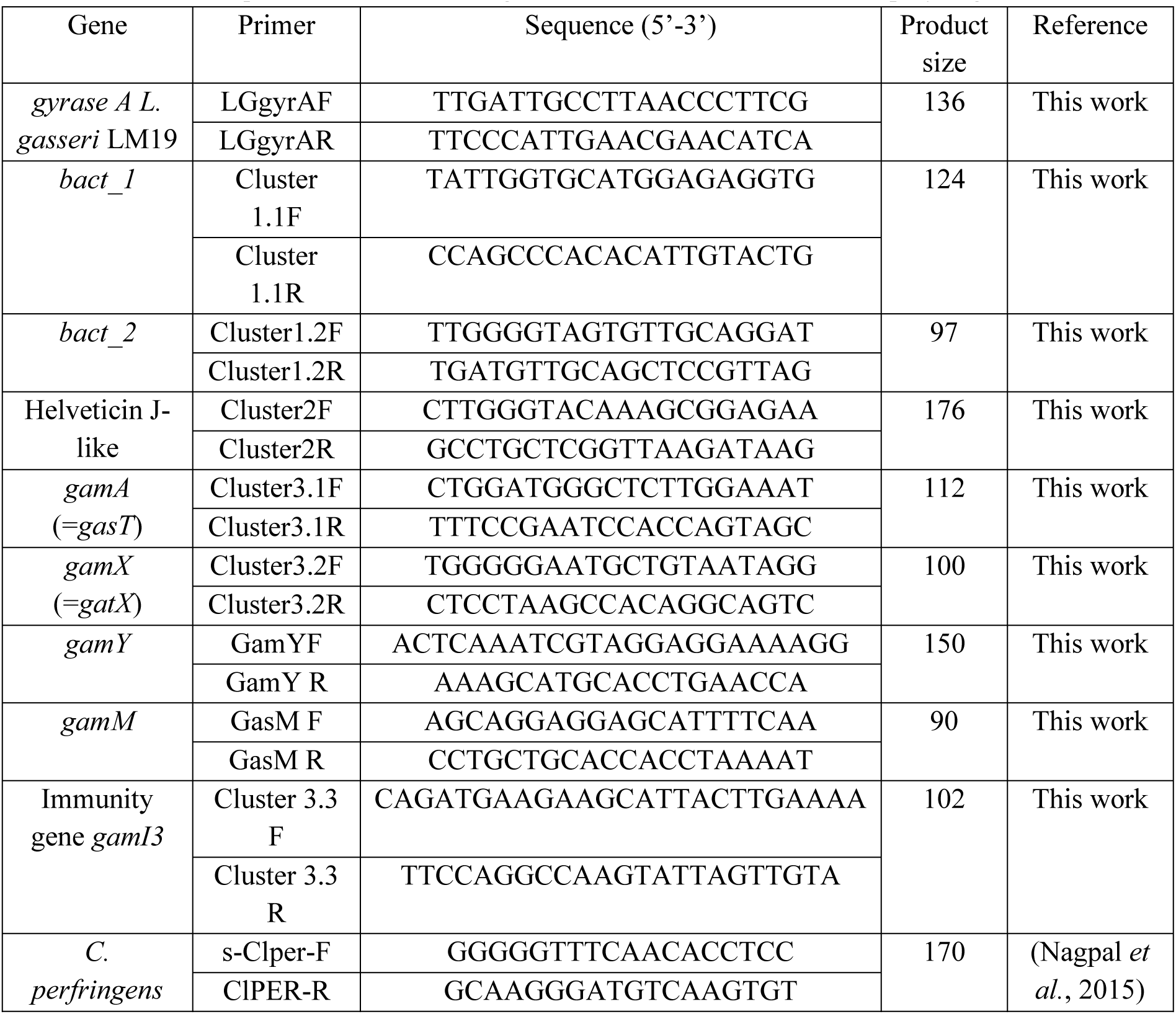
Primers for qRT-PCR studies in *L. gasseri* LM19 and detection of *C. perfringens*.

For DNA extraction, 16S rRNA amplification and sequencing and 16S rRNA-based metataxonomic analysis, 3 ml of each aliquot at different time points was treated for DNA extraction using the FastDNA Spin Kit for Soil (MP Biomedicals) following manufacturer’s guidelines. Total DNA concentration was measured by Qubit 3 (Invitrogen, UK) and normalized. The V4 region of the 16S rRNA gene was used for high throughput sequencing using the Illumina MiSeq platform. Data analysis was conducted using Quantitative Insights into Microbial Ecology (QIIME2 version 2018.11) (Bokulich *et al.*, 2018). A standard curve for *C. perfringens* NCTC 3110 was constructed by extracting gDNA as described previously (Ladero *et al.*, 2011) at different concentrations (colony forming units (cfu)/ml) of *C. perfringens* NCTC 3110. Each DNA concentration was measured using qPCR to determine the cycle signal associated with each cell concentration. Colon model treatments were analysed by RT-qPCR and total cfu were calculated for each treatment.

SCFA were measured using proton NMR (Parmanand *et al.*, 2019). Dilution buffer (D_2_O: 0.26 g NaH_2_PO_4_, 1.44 g K_2_HPO_4_, 17.1 mg sodium 3-(Trimethylsilyl)-propionate-d4 (TSP), 56 mg NaN_3_ in 100 ml) was mixed with cell-free sample supernatant in a proportion 1:10. 500 µl were collected in 5 mm NMR Tubes (GPE Scientific Ltd, UK). High resolution ^1^H NMR spectra were recorded on a 600 MHz Bruker Avance III HD spectrometer fitted with a 5 mm TCI cryoprobe and a 60 slot autosampler (Bruker, Germany). Sample temperature was controlled at 300 °K and the D_2_O signal was used as lock. Each spectrum consisted of 512 scans of TD = 65,536 data points with a spectral width of 20.49 ppm (acquisition time 2.67 s). The *noesygppr1d* presaturation sequence was used to suppress the residual water signal with low power selective irradiation at the water frequency during the recycle delay (D1 = 3 s) and mixing time (D8 = 0.01 s). A 90° pulse length of approximately 8.8 µs was used, with the exact pulse length determined automatically for each sample. Spectra were transformed with 0.3 Hz line broadening and zero filling and were automatically phased and referenced (to TSP) using the TOPSPIN 3.2 software. The resulting Bruker 1r files were converted to Chenomx (.cnx) format using the ‘Batch Import’ tool in the Processor module of Chenomx NMR Suite v8.12 with the TSP concentration set to 0.1 mM. Concentrations were obtained using the Chenomx Profiler module (Chenomx Inc, Canada).

### Statistical analysis

Significant differences between groups were established using a paired *t*-test, assuming normal distribution, equal variances. Both sides of the distribution were considered. Significance was considered when *P* value was <0.05.

## Results

### Antimicrobial activity

The antimicrobial activity of *L. gasseri* LM19, was assessed against a variety of Gram-positive and Gram-negative pathogens (Table 2), using a number of different approaches. The assay method affected the outcome, with the targets typically being more sensitive to LM19 grown on agar than to its cell free supernatant. The growth of *C. perfringens* NCTC 3110 and *L. bulgaricus* 5583 was inhibited by overlay assays. Cross-streaks showed antimicrobial activity against all Gram-positive indicators and *C. sakazakii*, while supernatants exhibited activity by filter disc against *L. bulgaricus*, *C. jejuni* and *M. luteus*. Well-diffusion assay only inhibited the growth of *L. bulgaricus*. As *L. bulgaricus* was the most sensitive indicator, it was used in subsequent tests.

**Table 2.**
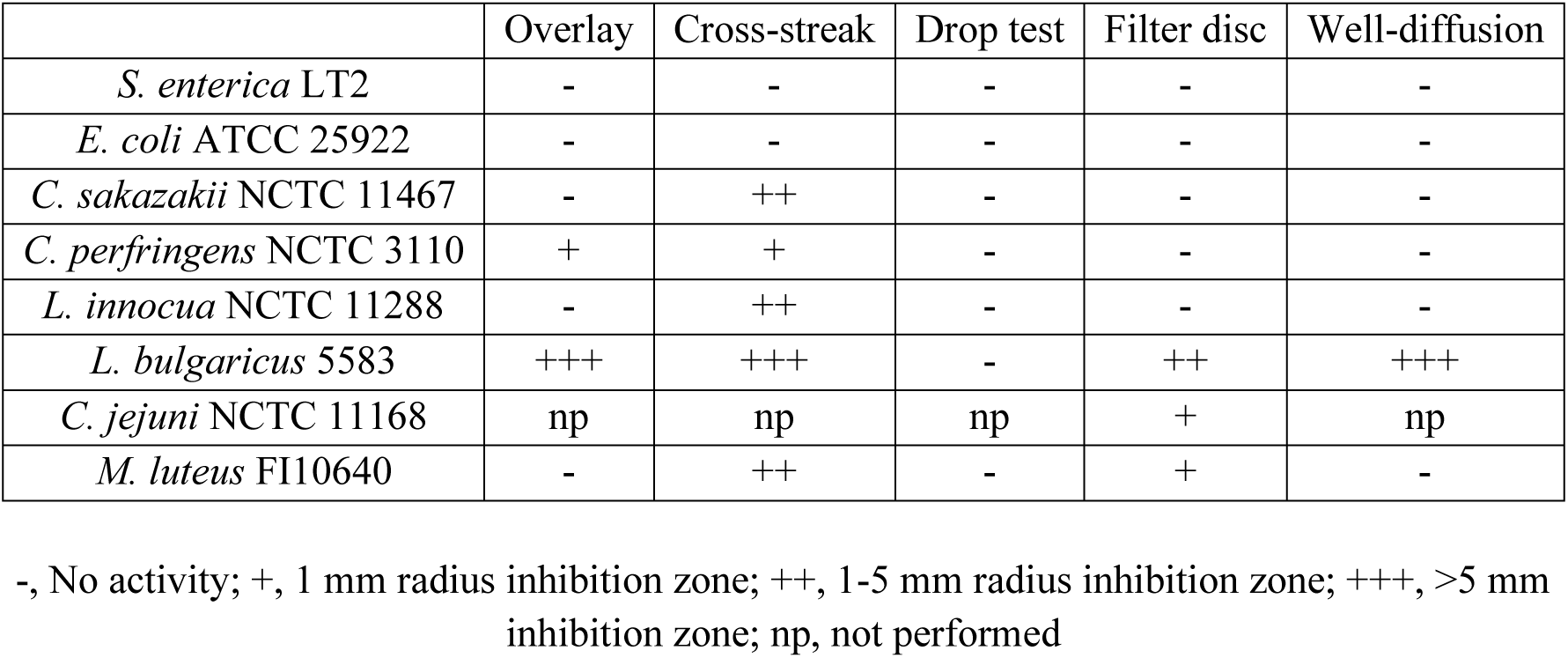
Summary of inhibitory activity of *L. gasseri* LM19 using different techniques

### Identification of bacteriocin gene clusters in the genome of *L. gasseri* LM19

The sequenced genome of *L. gasseri* LM19 was assembled into contigs and submitted to the NCBI under accession number SHO00000000. RAST analysis failed to reveal any bacteriocin clusters of interest, antiSMASH 3.0 indicated the presence of a single Microcin M-like cluster, while BAGEL 4, which specifically targets regions with bacteriocin similarities, found three clusters predicted to encode a number of potential bacteriocins. Manual investigation confirmed the presence of two clusters, whose putative structural peptides showed a high similarity to previously identified antimicrobial peptides from Class IIb bacteriocins (clusters 1 and 3), and a helveticin-like protein (cluster 2). The latter contained no other bacteriocin-associated genes on the basis of Blastp analysis; the product of the single gene showed 31.9 % identity and 43.1% amino acid consensus to helveticin J, which was originally characterised in *Lactobacillus helveticus* following heterologous expression (Joerger and Klaenhammer, 1986).

Cluster 1 (939 bp) is highly similar to the class IIb gassericin K7A cluster (EF392861) with 99% nucleotide identity. The cluster was predicted to encode two short peptides with leader sequences (Bact_1 and Bact_2) and a putative immunity protein (Fig.1a). Bact_1 and Bact_2 show 100% amino acid similarity with the gassericin S structural peptides GasA and GasX respectively (Kasuga *et al.*, 2019), while the surrounding genes do not resemble any other genes known to be associated with bacteriocin production. The putative immunity protein showed 97% amino acid homology to those of the acidocin LF221A and gassericin S clusters (Kasuga *et al.*, 2019; Majhenič *et al.*, 2004).

**Fig. 1.**
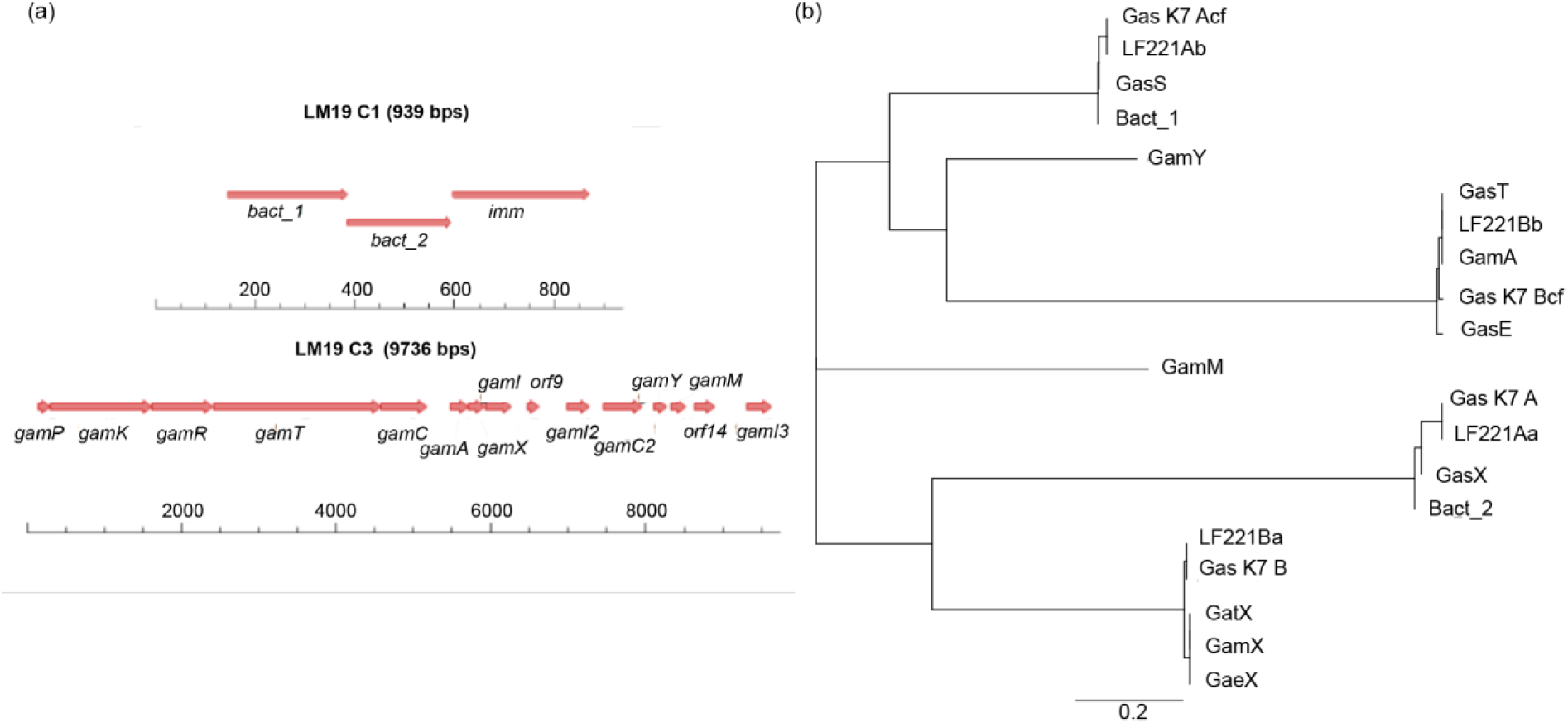
**a** Cluster LM19 C1, encoding Bact_1 and Bact_2, and Cluster LM19 C3 encoding GamA, GamX, GamY and GamM; **b** Phylogenetic tree of the amino acid sequences of putative bacteriocins identified in *L. gasseri* LM19 in context with the other class IIb gassericins.

Cluster 3 is 9736 bp in length and the first open reading frames (orfs) 1-8 show a high nucleotide homology to the gassericin T cluster from *L. gasseri* LA158 (AB710328, 99% over 100% coverage) and the gassericin E cluster from EV1461 (KR08485, 99% over 95% coverage) (Fig. 1a). There are two structural peptide-encoding genes, *gamA* and *gamX*, that are preceded by homologues of the gassericin E cluster as described previously (Maldonado-Barragán *et al.*, 2016). It is likely that they perform the same predicted functions as their gassericin E homologues, i.e., *gamP*, *gamK*, *gamR* for regulation, *gamT* and *gamC* for transport and, after the structural peptides, *gamI* for immunity, although a homologue to *gaeX* is missing. The predicted GamA peptide has the same sequence as GasT, Gas K7B cf and acidocin LF221B cf and has a single amino acid difference (W-L) from GasE (Table 3). The second putative peptide, GamX, has the same sequence as GatX and GaeX, all of which differ by a single amino acid (G-A) from Gas K7 B and acidocin LF221B (Table 3).

**Table 3.**
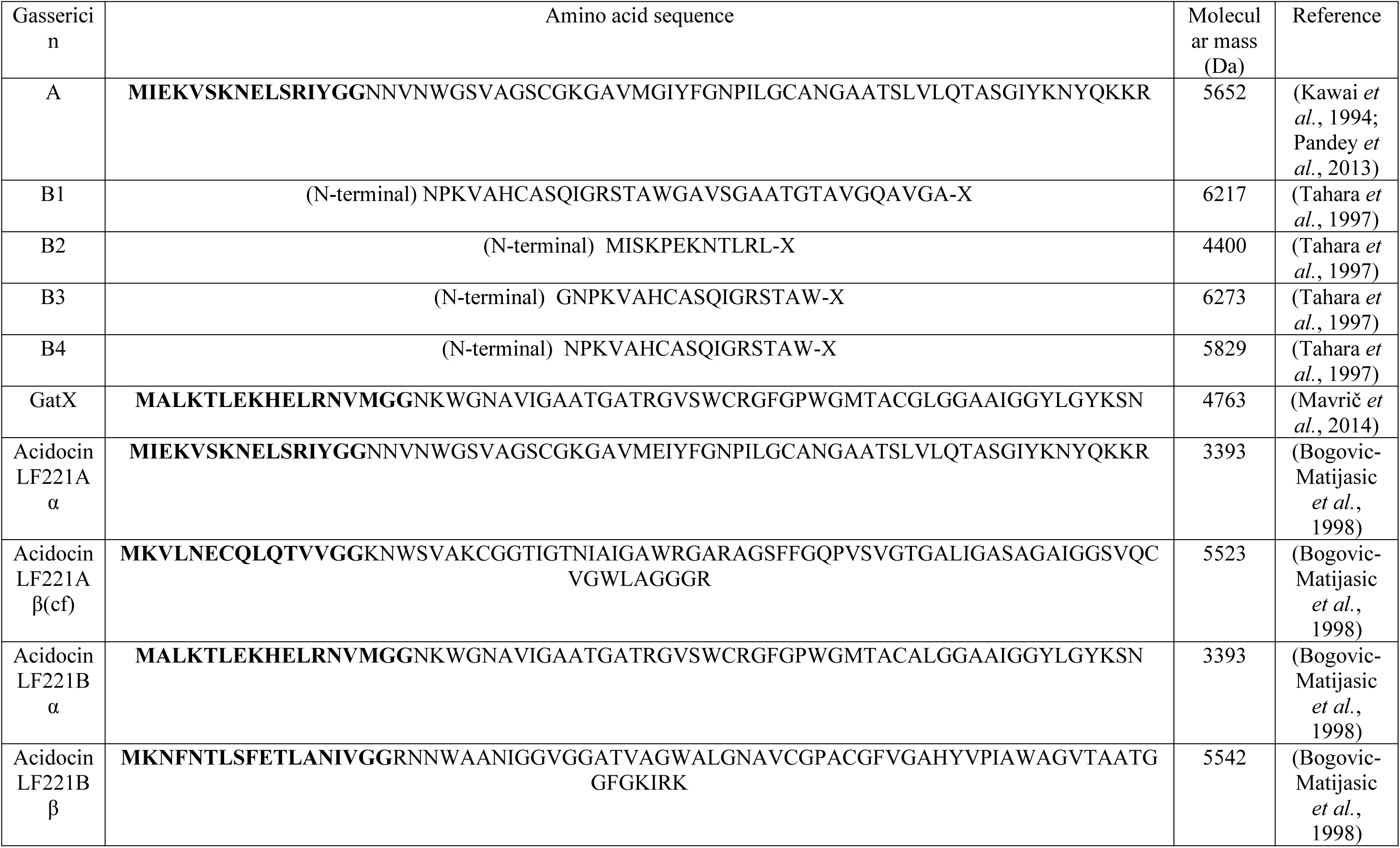
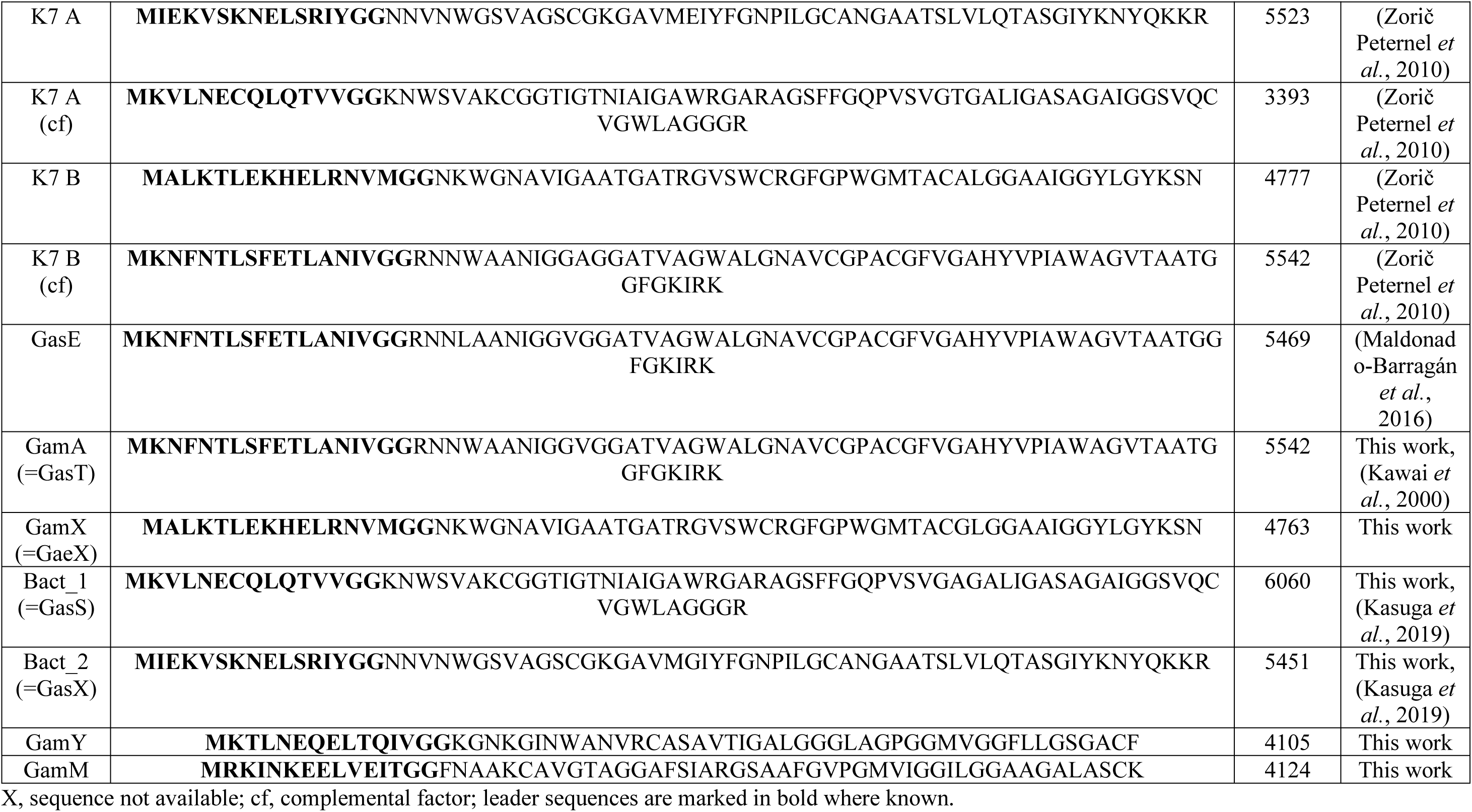
Bacteriocins described in *L. gasseri*.

In cluster 3, there are 7 further orfs including two additional putative structural genes, designated as *gamM* and *gamY*, which appear to encode a two-component bacteriocin. These putative peptides also show some similarity to other two-peptide component gassericins, but to a lesser extent than those previously reported (Fig. 1b). GamY shows similarity to GamM, with 25.4% identity and 47.6% consensus, and they both have similarity to K7 A cf (27.5% identity and 38.8% consensus; 25.3% identity, 44.3% consensus, respectively) and to GamA (18.7% identity, 33.3% consensus with GamM). Surrounding *gamY* and *gamM* are two genes encoding putative immunity proteins, GamI2 and GamI3, with homology to an enterocin A immunity domain (pfam 08951, 2.8e^−7^ and 1.1e^−6^, respectively), a putative transport accessory protein, GamC2, with some similarity to TIGR01295 bacteriocin transport accessory protein (1.18e^−9^), and thioredoxin superfamily cd02947 (5.21e^−7^), and two orfs with no matches. The genes on either side of the cluster show amino acid homology to transporters involved in solute or cation transport, and so are not predicted to be part of the cluster.

### Identification of antimicrobial peptides in culture

Cell and supernatant extracts from *L. gasseri* LM19 cultures were fractionated by HPLC and analysed by MS and their antimicrobial activity was assessed using *L. bulgaricus* 6901 as an indicator (Fig. 2 and 3). Antimicrobial activity was present in both cells and supernatant (Fig. 2a). MS shows that all peptide masses of interest are present.

**Fig. 2.**
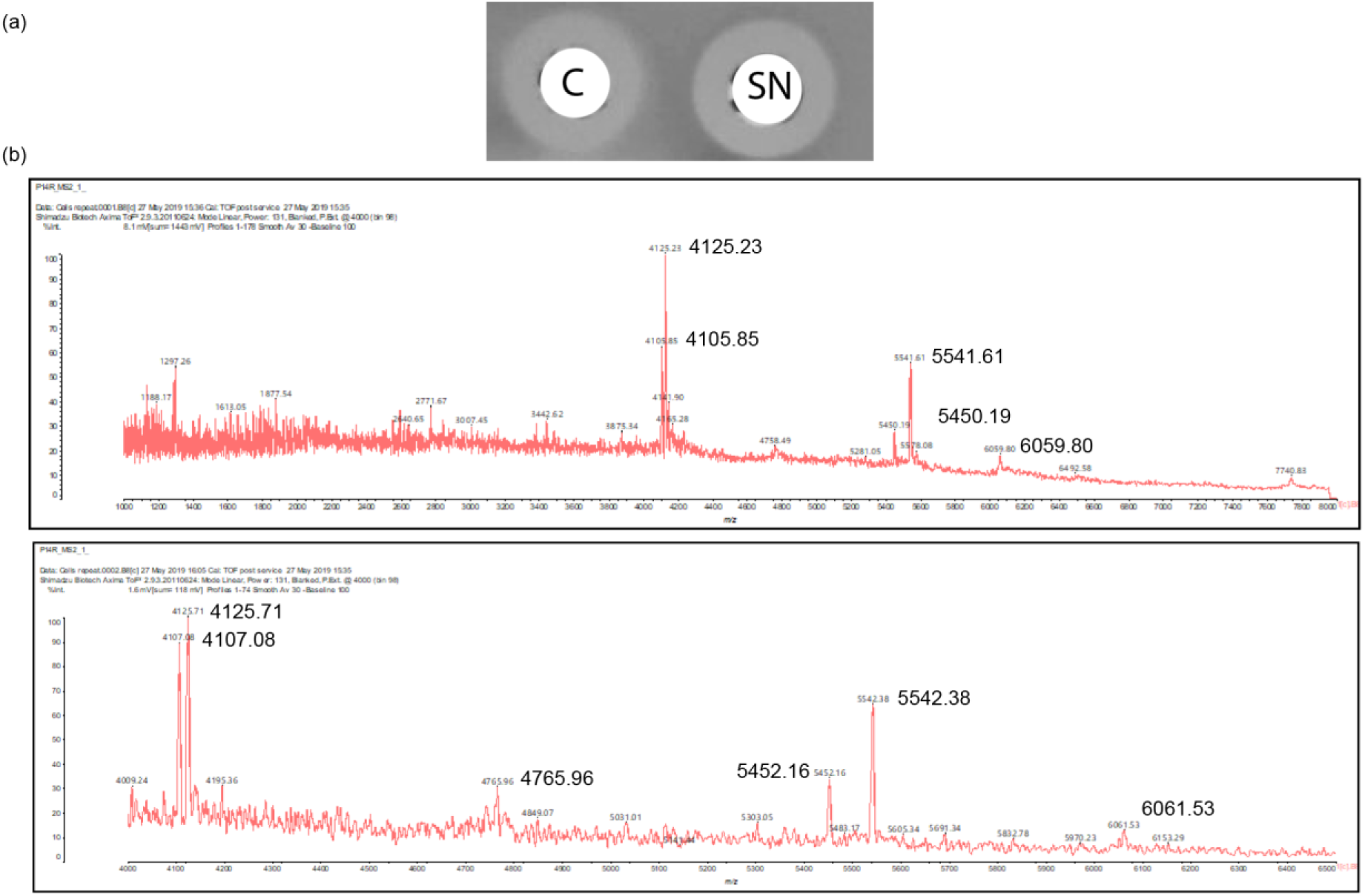
**a** Test for antimicrobial activity of cell (C) and supernatant (SN) fractions of *L. gasseri* LM19 culture; **b** Mass spectra showing peptide masses of interest in the cell extracts.

**Fig. 3.**
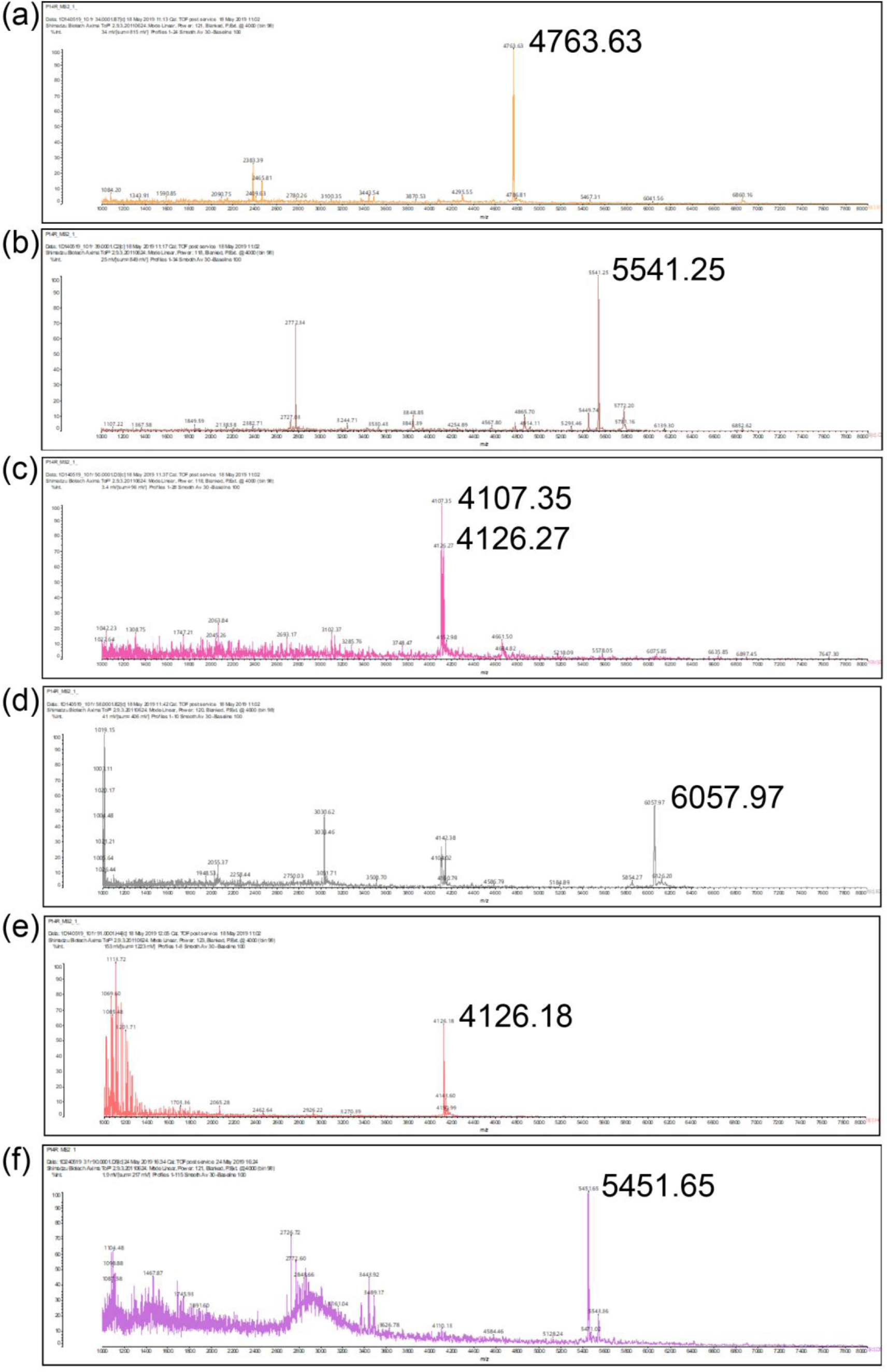
MS spectra of fractions showing putative masses for **a** GamX, 4763 Da; **b** GamA, 5541 Da; **c** GamM, 4126 Da and GamY, 4107 Da; **d** Bact_1, 6057 Da; **e** GamM, 4126 Da; **f** Bact_2, 5451 Da.

### Purification of bacteriocins from cells

MS analysis of HPLC fractions showed that many fractions contained one or more peptide masses that were consistent with those predicted by *in silico* analysis of the genome (Table 3, Fig. 2 and 3). mV response was very low (40 mV compared to around 1000 mV from a good producer), suggesting that production and consequently yield was very low. A mass corresponding to GamX (4763 Da) was detected in fractions 32-36 (Fig. 3a). Fractions 37-39 showed a mass corresponding to GamA, 5541 Da (Fig. 3b). Masses corresponding to GamY and GamM co-eluted in fraction 49-55 (Fig. 3c). Fractions 57-60 showed the expected mass from Bact_1, 6060 Da (Fig. 3d). Fractions 85-97 showed putative GamM mass, 4126 Da (Fig. 3e).

### Purification of bacteriocins from supernatants

MS analysis of the HPLC fractions 20-97 from the first HPLC run showed masses corresponding to Bact_1 and GamM (Fig. S1). Masses corresponding to putative Bact_1 eluted in fractions 59-65, while putative GamM mass was detected in fractions 76-97. Masses corresponding to Bact_2 and GamA eluted also in the first HPLC fractionation and were fractionated again. GamX eluted in fractions 86-90 (Fig. 3e), and also in fractions 66-68, co-eluting with GamA. Putative Bact_1 eluted in fractions 58-60. GamY eluted in fractions 69-72, and GamM showed in fractions 86-91 (Fig. S2 and S3).

### Synergy between peptides

Three sets of fractions were compared, fractions from HPLC run I, fractions from HPLC II run and synthetic peptides resuspended in milli Q water at 1 mg/ml (Fig. 4a). All synthetic peptides, except GamY, showed antimicrobial activity, with the highest activity coming from GamA and Bact_2. Fig. 4b and c show synergy assays between synthetic peptides. We observed clear synergy between Bact_1 and Bact_2 and a possible synergy between Bact_1 and GamA. No synergy was observed between GamM and GamY or between GamA and GamX.

**Fig. 4.**
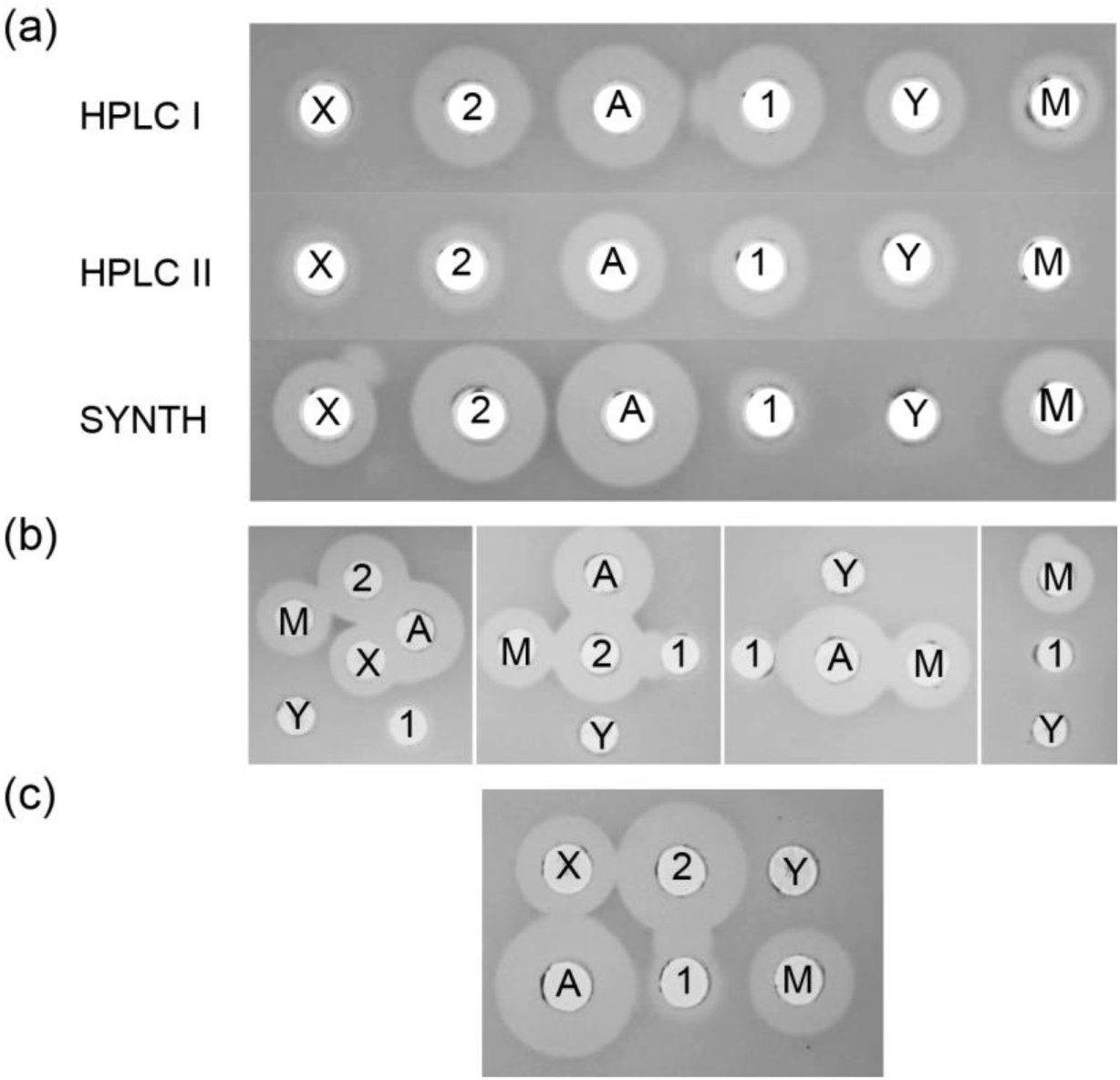
**a** Activity of GamX, (X); Bact_2, (2); GamA, (A); Bact_1, (1); GamY (Y); GamM, (M) from HPLC runs I and II containing putative peptide masses and synthetic peptides; **b** Synergy between the different peptides; **c** Synergy between pairs of peptides.

### Complex carbohydrates can favor viability and antimicrobial activity of *L. gasseri* LM19

*L. gasseri* LM19 was grown in colon model medium, simulating gut conditions, or home-made MRS, alone or supplemented with simple sugars (glucose, lactose and galactose) or complex polymers (inulin, starch and pectin). In general, more viable cells were recovered from MRS; growth on simple sugars was highest at 24 h but, at 48 h, complex carbohydrates gave higher counts (Fig. 5a). Interestingly, growth in the absence of a carbon source at 48 h was similar to that with simple sugars. On batch model medium, cell counts with glucose were lower than with all other treatments, and starch and pectin improved growth at 48 h. Antimicrobial activity from batch model medium with glucose was almost as high as that from MRS despite a ∼3 log difference in cfu (Fig. 5b). Glucose and galactose supplementation showed the highest antimicrobial activity at 24 h while, complex carbohydrates produced the highest activity after 48 h. At 48 h, higher levels of antimicrobial activity correlated with lower pH values and higher levels of cfu.

**Fig. 5.**
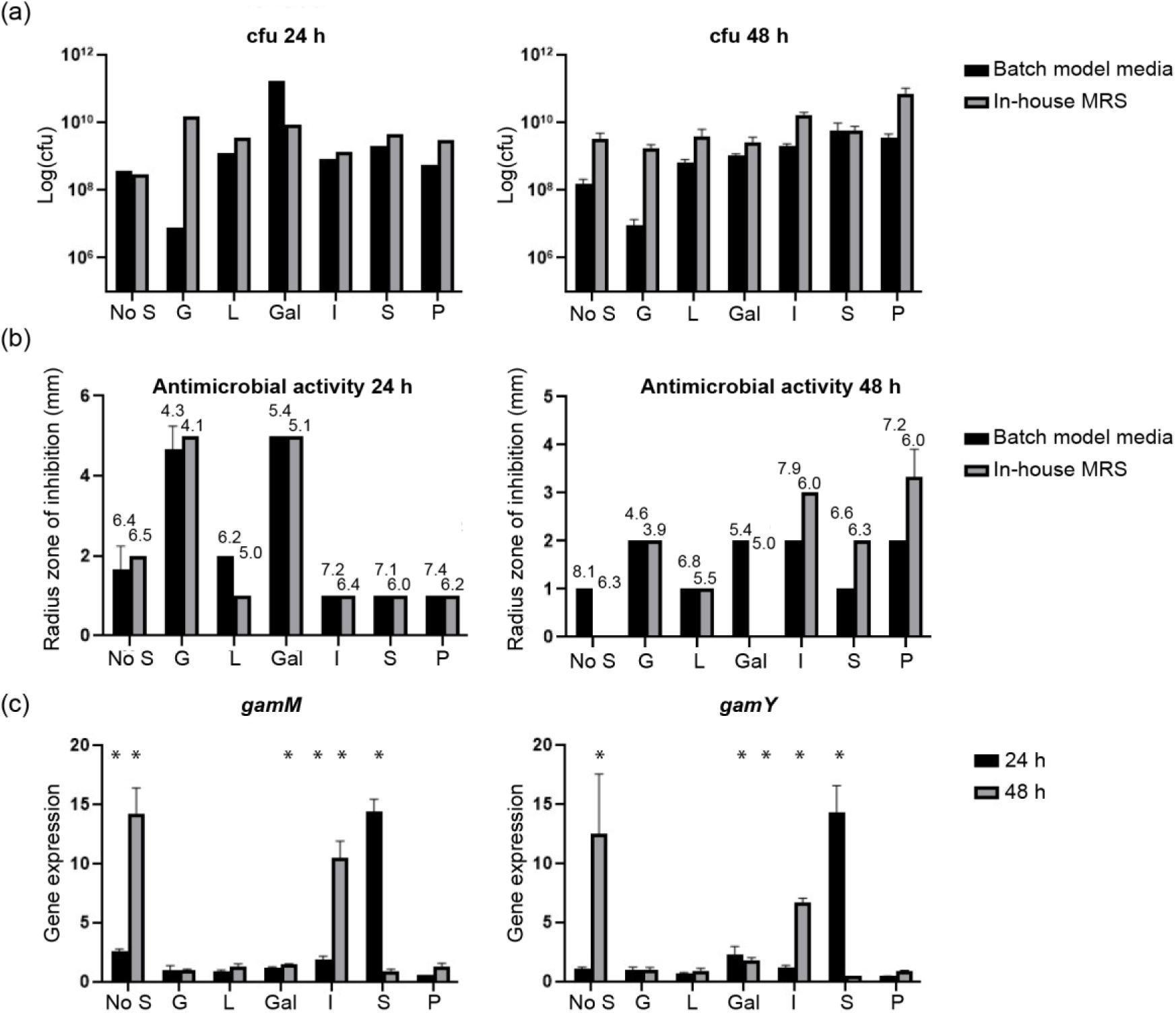
**a** Cfu of *L. gasseri* LM19 recovered when grown in batch model media or home-made MRS supplemented with different carbon sources; **b** Antimicrobial activity of cultures in **a** measured by well diffusion assay (Figures above bars indicate mean pH); **c** Gene expression levels of *gamM* and *gamY* after *L. gasseri* LM19 was cultured in home-made MRS supplemented with different carbon sources. No S, no supplementation; G, glucose; L, lactose; Gal, galactose; I, inulin; S, starch and P, pectin; *, significant difference to glucose supplementation (p<0.05). Results are the mean of triplicate measurements +/− standard deviation.

The changes in activity with carbon supplementation over time suggest control of antimicrobial production in different nutritional environments. Examination of bacteriocin gene expression in MRS also showed that no carbon supplementation increased the expression of *gamM* and *gam Y* significantly at 48 h. Starch supplementation increased the expression of both genes at 24 h, as did inulin at 48 h. Galactose supplementation also produced a significant increase in expression of *gamM* at 48 h and *gamY* at 24 and 48 h (Fig. 5c). Other bacteriocin genes did not show notable changes in expression, except for an increase in expression of the helveticin J-like gene in the presence of starch at 24 h (supplementary Table S1).

### *In vitro* colon model fermentations with *L. gasseri* LM19

Survival of *L. gasseri* LM19 and *C. perfringens* in an *in vitro* colon model *L. gasseri* LM19 was transformed with a plasmid conferring chloramphenicol resistance to allow selection and enumeration of this strain within a mixed microbial community. Transformation of electrocompetent cells gave an efficiency of 1.07×10^2^ transformants/ng of DNA. Fermentations with three different faecal donors were performed with four vessels per fermentation inoculated with *L. gasseri* LM19-pUK200, *C. perfringens* NCTC 3110, *L. gasseri* with *C. perfringens*, or a media control. *L. gasseri* numbers recovered increased from 5.3, 5.22 and 5.22 log_10_ cfu/ml in donors 1, 2 and 3, respectively at 4 h, to 6.12, 6.39 and 6.36 log_10_ cfu/ml at 8 h and 7.30, 7.31 and 7.47 log_10_ cfu/ml at 24 h. However, after 48 h, levels of recovery dropped to 3.66, 4 and 3.72 log_10_ cfu/ml.

*C. perfringens* levels were measured by qPCR, which detects DNA from both live and dead cells. Addition of *L. gasseri* LM19 did not have a negative effect on the *C. perfringens* population in the fermentation with faecal sample from donor 1; there was a tendency to lower *C. perfringens* counts in co-culture at 24 h with donors 2 and 3, but the changes were not significant (Fig. 6a).

**Fig. 6.**
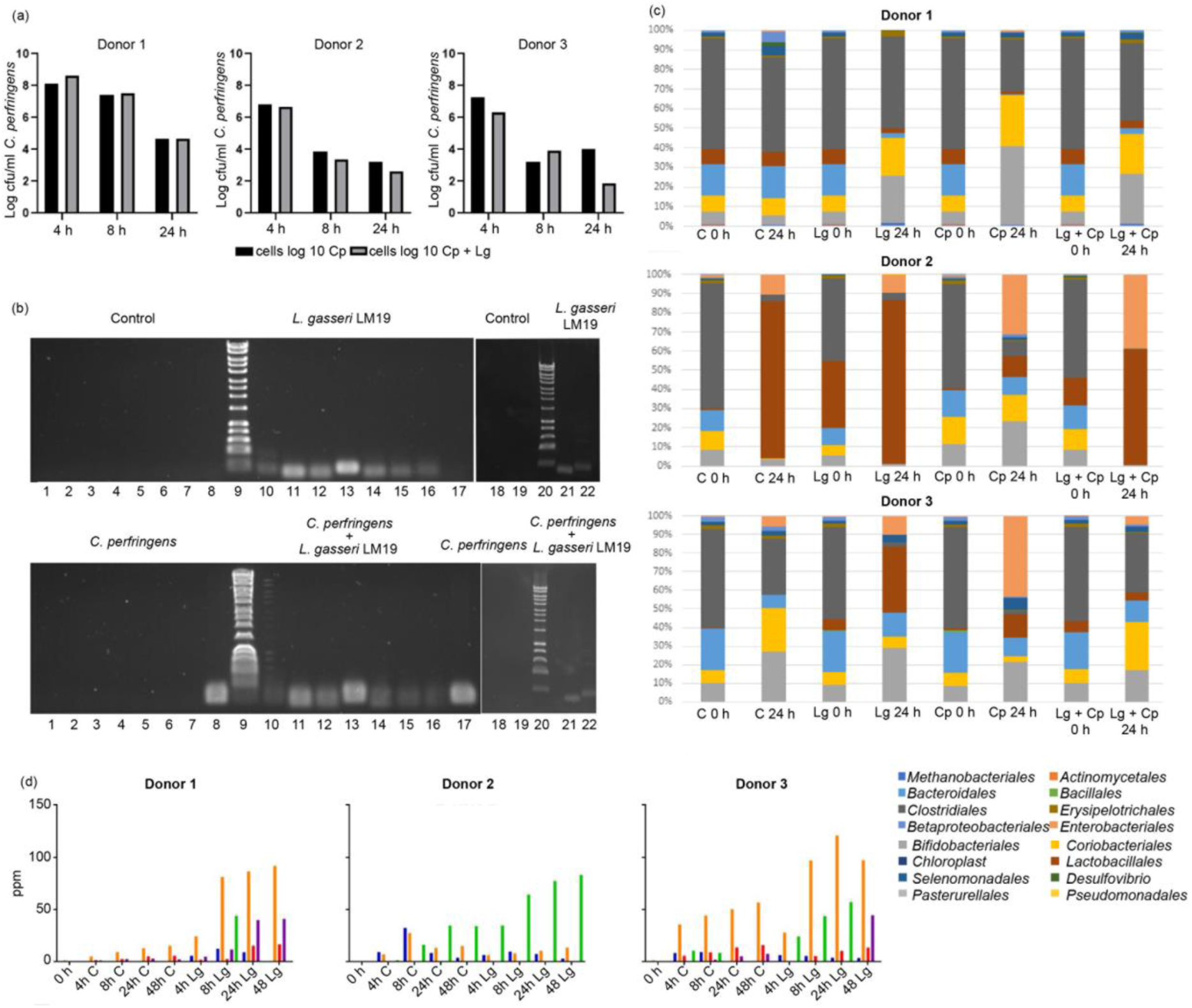
**a** *C. perfringens* NCTC 3110 population in the presence of *L. gasseri* LM19 in three different faecal fermentations; **b** Expression of bacteriocin genes in colon model at 24 h with donor 1; **c** Representation of relative abundance at order level in the faecal batch model fermentation for the three donors; **d** Production of SCFA in batch model faecal fermentation using inoculum from three different donors: blue, formate; orange, acetate; red, propionate; purple, butyrate; green, lactate.

#### Bacteriocin gene expression

PCR amplification of cDNA obtained from colon model samples using primers to detect the bacteriocin genes *bact_1, bact_2, helveticin-J like*, *gamA*, *gamX*, *gamM* and *gamY*, showed detectable levels of bacteriocin gene expression at 24 h (Fig. 6b). At 48 h, expression of only helveticin-J like, *gamM* and *gamY* genes was detected (data not shown).

#### Impact of *L. gasseri* LM19 on gut microbiota composition

Analysis of relative abundance at order, family and genus level was conducted. The initial bacterial composition was, as expected, different between the three donors (Fig. 6c). Bacterial populations from donor 1 remained relatively stable over 24 h. The addition of *L. gasseri* LM19, *C. perfringens* or both did not result in a significant increase in proportions of *Lactobacillales* or *Clostridiales*, and all 3 treatments resulted in similar increases in *Bifidobacteriales* and *Coriobacteriales* relative to the control, with the *L. gasseri* LM19 with *C. perfringens* treatment more similar to the *L. gasseri* LM19 only condition.

The initial population from donor 2 was constituted mainly of *Clostridiales*, with some *Bacteroidales, Coriobacteriales* and *Bifidobacteriales*. A change can be observed at 24 h in both the control and the samples where *L. gasseri* LM19 or *L. gasseri* LM19 with *C. perfringens* were added, with an increase in *Lactobacillales* along with a small increase in Enterobacteriales. The decrease in relative abundance of *Bifidobacteriales* and *Coriobacteriales* in the control, *L. gasseri* and *C. perfringens* + *L. gasseri* treatments was statistically significant (p<0.05) and not observed in the *C. perfringens* sample. *C. perfringens* alone appeared to prevent the overgrowth of *Lactobacillales*, while *Clostridiales* were decreased, being replaced by *Enterobacteriales*, *Bacteroidales*, *Coriobacteriales* and *Bifidobacteriales*. It was noted that addition of *L. gasseri* LM19 with *C. perfringens* gave a profile with more similarity to the control or *L. gasseri* LM19 samples.

In *L. gasseri* LM19 treatment of donor 3 samples, *Bifidobacteriales* and *Enterobacteriales* increased over time in a similar way to the control, but *Clostridiales* were almost completely replaced by *Lactobacillales*. This rise was not as large when the *L. gasseri* LM19 was co-inoculated with *C. perfringens*, while addition of *C. perfringens* alone did not manage to maintain levels of *Clostridales*, with increases seen in *Enterobacteriales*, *Bifidobacteriales* and *Lactobacillales*. In this case, the *L. gasseri* LM19 + *C. perfringens* treatment at 24 h was more similar to the control, with the exception of the presence of *Lactobacillales*, suggesting that the *C. perfringens* effect on the microbial composition was changed by the inoculation with *L. gasseri*.

#### SCFA analysis

Increases in the production of formic, acetic, propionic and butyric acids were observed in the three faecal fermentations in colon model conditions inoculated with *L. gasseri* LM9. However, there was a high variability in SCFA production between the three donors (Fig. 6d). In donor 1, production of SCFA, ethanol, succinate and, at 8 h only, lactate was increased compared to the control. In donor 2 there were notable increases in lactic acid from 4 h. Given the similar relative abundance of *Lactobacillales* (Fig. 6d) in control and *L. gasseri* treatment, this suggests an influence of *L. gasseri* LM19 on the native microbiota.

## Discussion

Here we report the ability of a representative of the human breast milk microbiota to exhibit antagonistic activity against different enteropathogens via production of previously identified bacteriocins and one novel bacteriocin. We have observed different carbon sources have an influence on the expression of these bacteriocin genes. *L. gasseri* LM19 survived and expressed these antimicrobial genes in a complex faecal environment under simulated colon conditions. This can be considered an important feature, since not all strains that exhibit probiotic traits are able to survive in colon conditions and, therefore, deliver their activity *in situ*. Additionally, we have observed other characteristics that are considered useful, such as the production of SCFA.

The presence of bacteria in human breast milk has been reported previously and the existence of a bacterial entero-mammary pathway has recently been proposed (Rodríguez, 2014). These bacteria might have a gut origin and that could explain their ability to survive in GI tract conditions and exhibit antagonistic traits against other gut bacteria such as enteropathogens that might share the same environment. Gassericins are antimicrobial peptides produced by *L. gasseri*. Several gassericins have been identified in sets of four, comprising two-peptide class II bacteriocins. K7 bacteriocins are a variant of acidocins LF221 and share similar sequences to GasT and its complementary peptide GatX, respectively, while GasE could be considered a variant of GasT. These two-peptide bacteriocins also show similarities with other two-peptide bacteriocins isolated from species previously grouped with *L. gasseri* (Tahara *et al.*, 1996). *L. gasseri* LM19 also presents two clusters of bacteriocins that show homology to acidocin LF221A and Gas K7A on one hand and acidocin LF221B and Gas K7B on the other hand. Additionally, we observed the presence of structural genes corresponding to a new two-component bacteriocin that show a greater variation in sequence to previously described gassericins. Partial purification of the products of these structural genes was conducted and we observed the presence of masses matching the expected size in eluted fractions that exhibited antimicrobial activity. Particularly, masses predicted to match those of the new potential peptides GamM (4124 Da) and GamY (4105 Da) were associated with antimicrobial activity. Synthetic peptides confirmed the activity of GamM but GamY did not show activity or synergy with GamM. Synergistic activity was reported previously between GasT and GatX and between the two components of gassericin S (here Bact_1 and Bact_2), respectively (Kasuga *et al.*, 2019). While Bact_1 and Bact_2 showed synergistic activity, the GasT and GatX homologues from this study, GamA and GamX, did not. However, we could observe a possible synergistic activity between GamA and Bact_1.

We demonstrated that *L. gasseri* LM19 is able to survive in simulated colon conditions within a complex faecal microbiota. Moreover, it is capable of expressing the bacteriocin genes in this environment. In previous work it was demonstrated that another potentially probiotic *L. gasseri*, strain K7, which produced 2 two-peptide bacteriocins K7 A, K7 A (cf), K7 B and K7 B (cf), was able to survive in faecal samples. Its bacteriocins were also the focus of examination by conventional PCR and RT-PCR (Treven *et al.*, 2013). In that instance the authors noted that bacteriocin genes were amplified by PCR from other LAB species present in the environment. However, in our controls and treatments where *L. gasseri* LM19 was not present, no PCR products were detected.

*L. gasseri* LM19 showed mixed effects on a strain of *C. perfringens* added to faecal fermentations of three different donors, causing a slight decrease in *C. perfringens* in only 2 out of 3 fermentations. This might indicate that the surrounding microbiota plays a synergistic or antagonistic role on the effect of *L. gasseri* LM19. However, it should be noted that in antimicrobial assays *C. perfringens* was only inhibited by *L. gasseri* LM19 cells, not cell-free supernatant, which might suggest that they should be in close proximity for an antimicrobial effect. However, co-inoculation of *L. gasseri* LM19 with *C. perfringens* did seem to alter the effect of *C. perfringens* on the background microbiota. In all three donors, the profiles seen after addition of *L. gasseri* LM19 with *C. perfringens* were more similar to instances where *L. gasseri* was added alone or control samples than to samples containing only *C. perfringens*.

Colon model fermentations also allowed the production of formic, acetic, propionic and butyric acids to be quantified using NMR. We observed an increase in SCFA production in the faecal fermentation of the three donors. However, as with the microbial composition, the amount of each SCFA produced was very different from one donor to another, which might be related to production by other members of the microbiota that varied between the three donors. In a previous study of consumption of a beverage prepared with *L. gasseri* CP2305, the stools of the participants presented an increased level of SCFA too, while the microbiota experienced some alterations, including an increase in the presence of bacteria from *Clostridium* cluster IV, known for producing higher amounts of SCFA (Sawada *et al.*, 2016). The authors of that study could not conclude if the increase of SCFA was due to the effect of *L. gasseri* or due to the proliferation of bacteria that produced more SCFA. SCFA production also depends on diet and availability of nutrients in the gastrointestinal tract as well as the resident microbiota (den Besten *et al.*, 2013; Holmes *et al.*, 2017).

This work shows the ability of *L. gasseri* LM19, a multi-bacteriocin breast milk isolate, to survive in colon conditions. Its ability to express different bacteriocin genes, including a novel gassericin M, under these conditions, makes it a candidate for further application studies.

## Acknowledgements

The authors would like to thank Dr. Lee Kellingray for training in the colon model system and Dr. Lizbeth Sayavedra, Dr. Maria Diaz-Garcia, Dr. Bhavika Parmanand and Saskia Neuert for training in QIIME2.

## Ethical statement

Faecal samples were provided by different donors, from a study approved by the QIB Human Research Governance committee (IFR01/2015) registered at http://www.clinicaltrials.gov (NCT02653001). The participants provided their written informed consent to participate in this study.

