## Supplementary material for "Production of multiple bacteriocins, including the novel bacteriocin gassericin M, by *Lactobacillus gasseri* LM19, a strain isolated from human milk": Table S1

Applied Microbiology and Biotechnology

Supplementary Information

Enriqueta Garcia-Gutierrez[1,2] 0000-0001-5683-7924
Paula M. O'Connor[2,3] 0000-0001-6462-2077
Ian J. Colquhoun[1]
Natalia M.Vior[4] 0000-0003-1890-3884
Juan Miguel Rodríguez[5]
Melinda J. Mayer[1] 0000-0002-8764-2836
Paul D. Cotter[2,3*] 0000-0002-5465-9068
Arjan Narbad[1]

[1] Gut Microbes and Health, Quadram Institute Bioscience, Norwich, UK

[2] Food Bioscience Department Teagasc Food Research Centre, Moorepark, Fermoy, Cork, Ireland

[3] APC Microbiome Ireland, University College Cork, Cork, Ireland

[4] Molecular Microbiology, John Innes Centre, Norwich, UK

[5] Dpt. Nutrition and Food Science, Complutense University of Madrid, Madrid, Spain

*Corresponding author:

Paul D. Cotter

**Table S1** Expression levels of the different bacteriocin genes when *L. gasseri* LM19 is grown in MRS supplemented with different carbon sources.

| | | No supplemental carbon source | Glucose | Lactose | Galactose | Inulin | Starch | Pectin |
|---|---|---|---|---|---|---|---|---|
| *Helveticin J-like* | 24 h | 2.214493 | 0.813366 | 0.378005 | 0.945059 | 2.103041 | 6.092256 | 3.206449 |
| | | 2.984523 | 0.940485 | 0.896874 | 0.879025 | 2.131659 | 8.233333 | 2.338327 |
| | | 3.140462 | 1.246149 | 1.791883 | 1.420084 | 2.3497 | 4.583586 | 3.460491 |
| | 48 h | 1.533635 | 1.113387 | 2.123516 | 0.923994 | 3.169939 | 2.619797 | 4.684695 |
| | | 2.015943 | 1.207862 | 1.918469 | 0.539597 | 1.804945 | 2.331818 | 1.708174 |
| | | 1.785436 | 0.678751 | 2.512207 | 1.0777 | 0.795788 | 2.456814 | 3.552797 |
| *gamA* | 24 h | 3.189112 | 0.75088 | 1.017589 | 1.64793 | 0.118147 | 1.37712 | 0.216758 |
| | | ND | 0.570641 | 1.064853 | 2.266011 | 0.0836 | 0.712595 | 0.387201 |
| | | 1.552028 | 1.678479 | 0.549486 | 1.162842 | 0.097204 | 1.549341 | 0.323908 |
| | 48 h | 0.873485 | 1.03158 | 0.632815 | 1.059112 | 0.319271 | 0.134377 | 0.394844 |
| | | 1.117175 | 1.157374 | 0.454659 | 1.496254 | 0.657633 | 0.789141 | 0.473473 |
| | | 0.479252 | 0.811046 | 0.597229 | 0.882003 | 0.391302 | ND | 0.453086 |
| *gamX* | 24 h | ND | 0.806815 | 0.444671 | 0.434314 | 0.213722 | 0.707014 | 0.391155 |
| | | 3.625828 | 0.785025 | 0.901445 | 2.118159 | 0.184259 | 0.622787 | 0.349972 |
| | | 1.86591 | 1.408159 | 0.522068 | 1.340072 | 0.138294 | 2.219442 | 0.330979 |
| | 48 h | 1.124289 | 1.042114 | 0.786497 | 1.139194 | 0.837124 | 1.260515 | 0.524863 |
| | | 1.058861 | 1.332849 | 0.58217 | 0.935632 | 1.157902 | 0.802189 | 0.456603 |
| | | 1.139984 | 0.625037 | 0.708094 | 0.429733 | 0.818477 | ND | 0.51424 |
| *bact_1* | 24 h | 0.005805 | 0.404422 | 1.092349 | 1.805544 | 0.581332 | 0.018658 | 0.201163 |
| | | 0.002988 | 0.972273 | 0.813342 | 1.697526 | 0.802422 | 0.035524 | 0.299356 |
| | | 0.957559 | 1.623305 | 0.997877 | 0.990641 | 0.937203 | 0.070386 | 0.828994 |
| | 48 h | 0.032678 | 1.093899 | 1.150276 | 1.976544 | 0.823009 | 0.002208 | 0.809992 |
| | | 0.028547 | 0.751831 | 1.47988 | 2.044829 | 0.805792 | 0.001814 | 0.821299 |
| | | 0.025925 | 1.15427 | 1.092763 | 1.014297 | 0.988272 | 0.004044 | 0.699541 |
| *bact_2* | 24 h | 0.017065 | 0.574438 | 0.715599 | 0.659399 | 0.427271 | 0.06301 | 0.130917 |
| | | ND | 0.385345 | 0.682889 | 1.246362 | 0.317917 | 0.01852 | 0.203024 |
| | | 0.296936 | 2.040217 | 1.344177 | 1.201405 | 0.699416 | 0.042298 | 0.685973 |
| | 48 h | 0.049096 | 1.006778 | 1.435194 | 2.156581 | 0.550855 | ND | 0.949504 |
| | | 0.030485 | 0.800096 | 2.121002 | 1.312435 | 0.632109 | 0.016761 | 1.007825 |
| | | 0.039685 | 1.193125 | 1.780455 | 0.773914 | 0.596356 | 0.009461 | 0.058042 |

ND, not detected

HPLC I

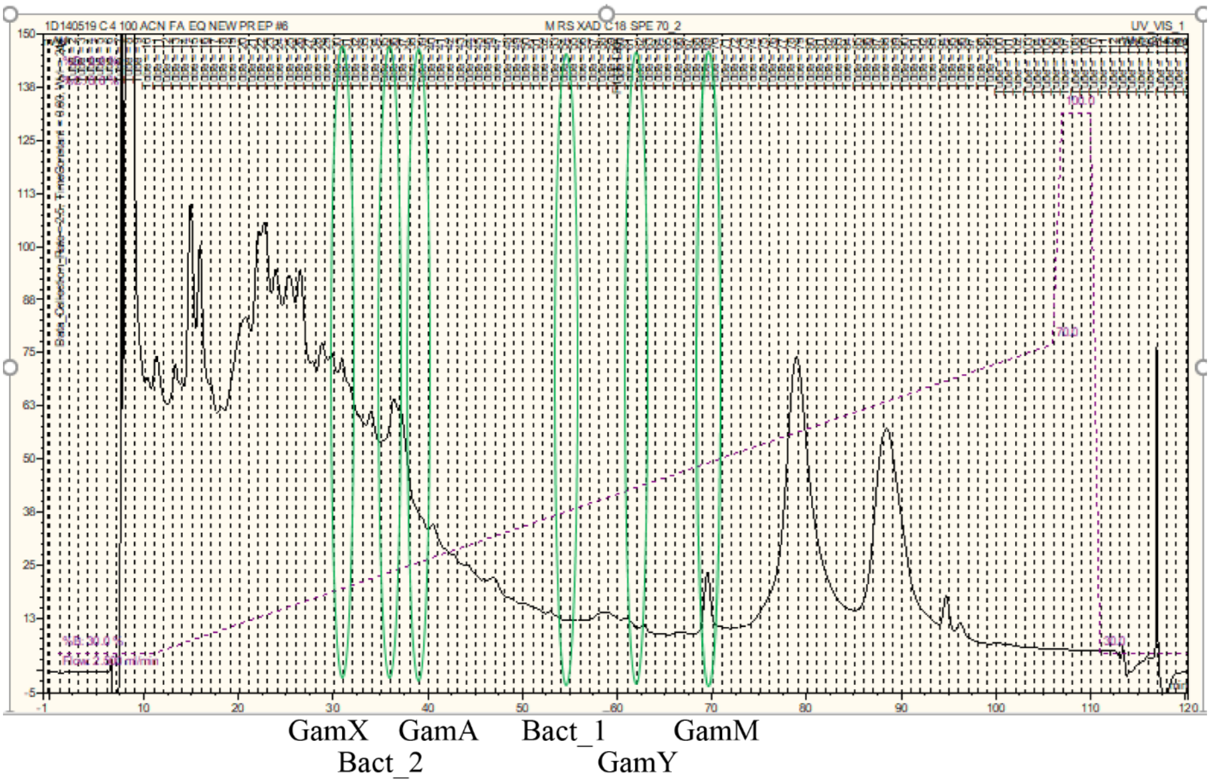


**Fig S1** HPLC elution fractions from *L. gasseri* LM19 active supernatant. MS chromatograms of fractions are shown in Figure S3.

HPLC II

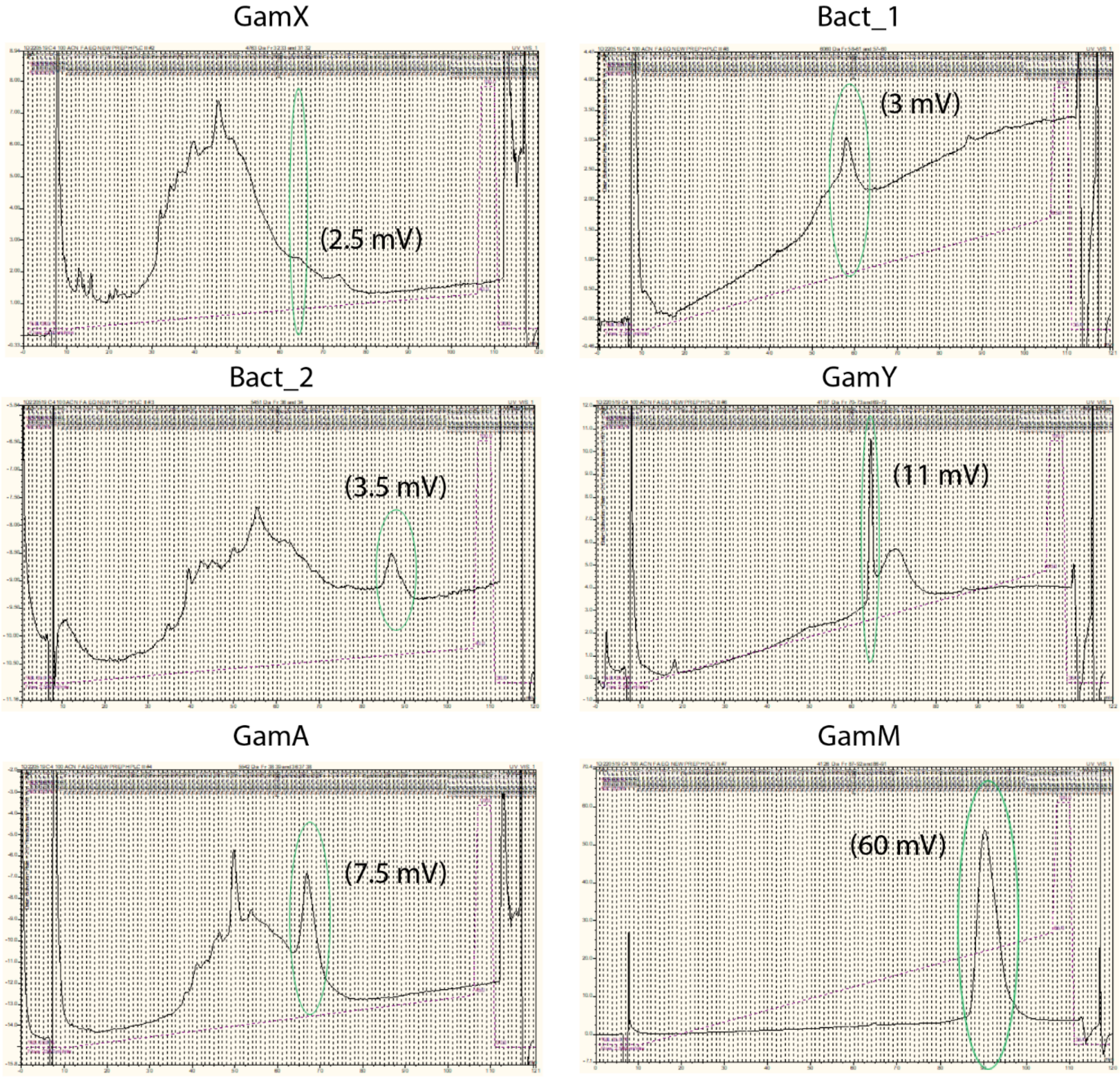


**Fig S2** mV levels of HPLC run of each HPLC I fraction that showed putative bacteriocin mass.

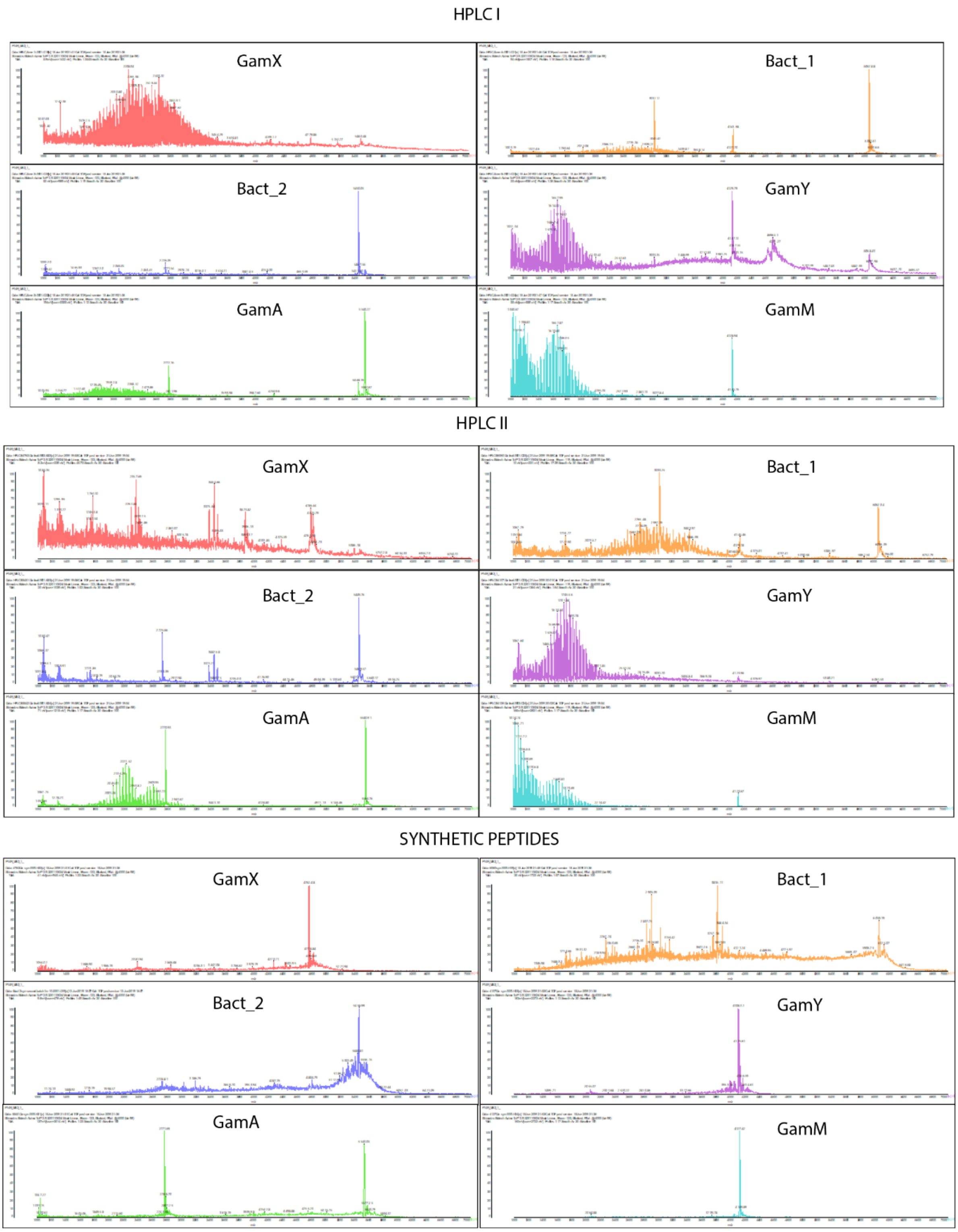


**Fig S3** MS chromatograms of assayed fractions from HPLC I, HPLC II and synthetic peptides.
